# Macrosystem community assembly patterns are predicted by foundation tree species genetic connectivity and environment across the American Southwest

**DOI:** 10.1101/2021.06.24.449837

**Authors:** Helen M. Bothwell, Arthur R. Keith, Julia B. Hull, Hillary F. Cooper, Lela V. Andrews, Christian Wehenkel, Kevin R. Hultine, Catherine A. Gehring, Samuel A. Cushman, Thomas G. Whitham, Gerard J. Allan

**Affiliations:** Environmental Genetics & Genomics Facility, Department of Biological Sciences, Northern Arizona University, Flagstaff, AZ 86011, USA; Research School of Biology, Australian National University, Canberra, ACT 2602, Australia; Tecan Genomics, Inc., 900 Chesapeake Drive, Redwood City, CA 94063, USA; Universidad Juárez del Estado de Durango, Durango 34120, México; Department of Research, Conservation and Collections, Desert Botanical Garden, Phoenix, AZ 85008, USA; Merriam-Powell Center for Environmental Research, Northern Arizona University, Flagstaff, AZ 86011, USA; Rocky Mountain Research Station, United States Forest Service, 2500 S. Pine Knoll Dr., Flagstaff, AZ 86001, USA

**Keywords:** biodiversity, community genetics, ecotype, macrosystems ecology, multiscale, *Populus fremontii*

## Abstract

Macrosystems ecology is an emerging science that aims to integrate traditionally distinct disciplines to predict how hierarchical interacting processes influence the emergence of complex patterns across local to regional and global scales. Despite increased focus on cross-scale relationships and cross-disciplinary integration, few macroecology studies incorporate genetic-based processes. Here we used a community genetics approach to investigate the pattern-process relationships underlying the emergence of macroscale biodiversity patterns. We tested the hypothesis that environmental variation, geography, and genetic connectivity in a foundation tree species differentially predict associated community assembly patterns from local to continental scales. Using genome-wide SNP data, we assessed genetic connectivity as a function of genetic similarity and structure in Fremont cottonwood (*Populus fremontii*) across its distribution throughout the southwestern US and México. For the same trees, we measured community composition, diversity, and abundance of leaf modifying arthropods and sequenced targeted amplicons of twig fungal endophytes. Five key findings emerged. (1) We identified three primary and six secondary population genetic groups within *P. fremontii*, which occupy distinct climate niches. (2) Both the leaf modifying arthropod and fungal endophyte communities were significantly differentiated across host tree ecotypes, with genetic distance among sampling locations explaining 13-17% of respective macroscale community structure. (3) For arthropods, environmental distance was the strongest driver of community similarity. (4) Conversely, host genetic connectivity was the most important contributor to macroscale endophyte community structure, with no significant contribution of environmental distance. (5) Furthermore, we observed a shift in the strength of interspecific relationships, with host genetics most strongly influencing associated communities at the intermediate population scale. Our findings suggest that genetic connectivity and environmental variation play integrated roles in macroscale community assembly, and their relative importance changes with scale. Thus, conservation genetic management of the diversity harbored within foundation species is vital for sustaining associated regional biodiversity.

## Introduction

Macroecological patterns arise from complex hierarchical interactions between biotic (e.g., gene flow, species interactions) and abiotic (e.g., climate, geography) factors operating across local to continental scales. Predicting macrosystem properties (e.g., biodiversity, community structure) requires understanding the underlying factors that drive pattern-process relationships across scales (Peters et al. 2007, 2008, 2011), and is a pressing challenge for mitigating widespread biodiversity loss under rapid global change (Carpenter 2006, Macdonald et al. 2020). Despite recent increased focus on cross-scale and cross-disciplinary integration (Heffernan et al. 2014, Fei et al. 2016, Dodds et al. 2021), there is a paucity of studies linking genetic-based processes to ecological patterns at large geographic scales (McGill et al. 2019).

Macrosystem properties are, in part, defined by evolutionary processes. At the scale of individuals, fine-scale processes (e.g., natural selection) influence the abundance and distribution of genetic variants across heterogeneous environments. At intermediate scales, population-level processes (e.g., dispersal, mating, genetic drift) contribute to the evolution of differentially adapted populations interacting across a regional landscape matrix. At broad scales, metapopulation dynamics are influenced by global climate patterns (e.g., post-glacial migration) resulting in the emergence of continental-scale phylogeographic structure. Within a community context, interspecific interactions are in part dictated by differential phenotypic expression, which has manifested as a result of individual- and population-level processes operating within species. Macrosystem properties are then the culmination of intraspecific evolutionary processes coupled with interspecific interactions among members of ecological communities, within the context of environmental and geographic factors driving pattern-process relationships across all scales. Research in the field of community genetics, which investigates the role of genetic-based interactions in the emergence of community properties, has the potential to advance macrosystems ecology by providing a theoretical framework for linking fine- and intermediate-scale genetic processes to landscape-level patterns (Mitton 2003, Agrawal 2003, Shuster et al. 2006, Allan et al. 2012, Whitham et al. 2006, 2020).

A critical question in macrosystems ecology is whether local pattern-process relationships can be scaled up to predict regional or continental scale patterns (Bangert et al. 2008). Omernik (1987) originally identified ecoregions as important units for managing broadscale biodiversity, a practice which has been widely adopted for guiding scientific studies and ecosystem management by state, federal, and non-governmental agencies (Omernik and Griffith 2014). Ecoregions, defined as large areas with similar climatic, biotic, and geophysical properties, can serve as useful proxies for managing regional biodiversity when detailed metapopulation data is lacking; yet, how well ecoregions define organizational structure varies among community members. For example, plants and vertebrates typically align more strongly with ecoregional boundaries than do arthropods and fungi (Smith et al. 2018). Identifying the abiotic and biotic processes that give rise to community composition and structure and determining how these factors differentially affect community assembly across spatial and temporal scales is critical to gaining a deeper understanding of macroscale biodiversity patterns (Mittlebach and Schemske 2015).

Similar to the concept of ecoregions, ecotypes denote populations and associated communities that form under the combined influence of shared biotic and abiotic factors (Turresson 1922, Hufford and Mazer 2003). Ecotypes are recognized as important units of local adaptation in forest trees (Cushman et al. 2014, Ikeda et al. 2017, Cooper et al. 2019, Bothwell et al. 2020, Blasini et al. 2021), shrubs (Yang et al. 2015, Germino et al. 2019), grasses (Lovell et al. 2021), insects (Raszick and Song 2016) and fungi (Leonhardt et al. 2019). In the case of foundation species (Ellison et al. 2005), host plants represent a biotic environment that provides critical habitat and modulates ecosystem services that a wealth of other species depend on (Lamit et al. 2015, Keith et al. 2017). Genetic variation underlying plant phenotypic variation (e.g., phytochemistry, phenology) extends the influence of host plants, resulting in predictable community and ecosystem phenotypes (Whitham et al. 2006, 2020).

The positive correlation between plant genetic similarity and associated community similarity (Barbour et al. 2009, Zytynska et al. 2012), described by Bangert et al.’s “Genetic Similarity Rule” (2006), highlights an important link between plant genetics and the emergence of higher-level community structure. Plant genotype is a known driver of community diversity and structure at local scales (e.g., common garden; Ferrier et al. 2012), however, some evidence suggests environmental variation may have a stronger influence at larger spatial scales (Johnson and Agrawal 2005, Bangert et al. 2008, Busby et al. 2014). For example, Bangert et al. (2006) found the relationship between tree genetic similarity and community similarity decreased as the scale of investigation increased from stands to river systems to regions. While previous studies have explored this relationship at the species level, we extend the framework here to include genomic data for both a foundation tree host and its associated community.

Here we investigated how genetic connectivity in a foundation tree species, *Populus fremontii*, influences local to continental-scale community assembly across southwestern North America. We hypothesized that (1) host tree genetic variation, geography, and environment would differentially predict community similarity. We investigated these relationships for two distinct communities: leaf modifying arthropods and twig fungal endophytes. These assemblages directly interact with plant tissues, cuing into growth rates, phenological, morphological, physiological, and phytochemical variation, all of which are under genetic control in this system (Rehill et al. 2005, Grady et al. 2013, Fischer et al. 2017). For example, fungal endophyte abundance has been shown to vary with *P. fremontii* condensed tannins (Bailey et al. 2005). Leaf modifying arthropods interact with leaves via gall formation, herbivory, and altering the physical structure of leaves (e.g., rolling/tying leaves to form protective enclosures); conversely, endophytes exist fully embedded within the host’s cellular matrix. Given the more intimate association of fungal endophytes living within the host’s tissue, (2) we predicted that tree genotype would be a stronger predictor for endophytes compared to arthropods. We further predicted that (3) plant genetic similarity would be a stronger predictor of community composition and structure at the local scale, whereas geographic and environmental distance would exert a more prominent influence on community similarity at broad scales. Tests of these hypotheses will help integrate community genetics theory with macrosystems ecology, providing an evolutionary basis for understanding landscape-level patterns of community assembly, and thus improve our capacity to support effective conservation policy and biodiversity management at regional to continental scales (Heffernan et al. 2014).

## Methods

### Species & collection information

Fremont cottonwood (*Populus fremontii*) is a foundation riparian tree that occurs throughout the southwestern US and northwestern México. Previous research has identified three *P. fremontii* ecotypes within the coterminous US, based on population genetic structure and environmental niche differentiation (Cushman et al. 2014, Ikeda et al. 2017). We collected leaf material and geographic coordinates for 453 trees at 58 sampling locations, stratified to maximize geographic and climatic variation across the three ecoregions (Table S1).

### Community data

Leaf modifying arthropod and twig fungal endophyte communities were sampled on a subset of the trees collected above during the summer of 2014 (late May to early August). Leaf modifying arthropods (e.g., galling insects, leaf tiers, leaf rollers, leaf miners) create distinctive, species-specific structures that conveniently allow for identifying species in their absence and reduce effects of temporal turnover when sampling across broad geographic regions (Bangert et al. 2006). Temporal bias was further minimized by conducting surveys from south to north, following clines in host tree phenology. For example, *P. fremontii* genotypes collected from southern Arizona to central Utah and grown in a common garden exhibited substantial variation in timing of spring bud flush, spanning a range of 55 days (Cooper et al. 2019).

Arthropod surveys were standardized by branch diameter (2-3 branches/tree, ~35mm total) to account for leaf area, and survey time (15min/tree). Arthropod species were visually identified to the lowest possible taxonomic level; any unrecognized species were collected, later identified in the lab, and added to the Northern Arizona University Insect Collection. To assess twig fungal endophyte communities, we collected 10 twigs/tree, including 3-years growth, directionally stratified around each tree’s circumference. Twigs were frozen prior to genomic sequencing, and variation in growth characteristics quantified among ecoregions.

### Plant genetic data

To assess population genetic structure and connectivity, we selected 5-6 trees per sampling location for genomic analysis. Leaf material was dried in Dri-Rite©, and ~0.2g per sample was ground with a 2000 Geno/Grinder (SPEX, SamplePrep, Metuchen, NJ, USA). Genomic DNA was extracted with DNeasy 96 Plant Mini Kits (Qiagen, Valencia, CA, USA) and quantified using Quant-iT PicoGreen on a Synergy HTX Microplate Reader (BioTek Instruments, Inc., Winooski, VT, USA). DNA was standardized to 5ng/μL, and double digest restriction-associated DNA (ddRAD) libraries were prepared following a modified Peterson et al. (2012) protocol. Briefly, 25ng DNA was digested with MspI and EcoRI restriction endonucleases and ligated to double stranded adapters in 20μL reactions. Ligation products were amplified with indexed primers for 15 PCR cycles. Indexed ligation products were purified with PEG-8000 and Sera-Mag Speedbeads Carboxylate-Modified Particles (Thermo Scientific, Fremont, CA; Rohland & Reich, 2012). Indexed samples were then pooled and size selected for 200-350bp fragments using a Pippin Prep (Sage Science, Inc., Beverly, MA). Fragment size distribution was assessed using Bioanalyzer high sensitivity DNA chips (Agilent Technologies, Santa Clara, CA, USA), and final DNA concentrations were quantified via qPCR (Eppendorf Mastercycler Realplex 4; Eppendorf, Inc., Westbury, NY). Sequencing was performed on a MiSeq Desktop Sequencer Illumina, Inc. San Diego, CA) in 1×150 mode at Northern Arizona University’s Environmental Genetics and Genomics Laboratory (Flagstaff, AZ, USA).

Quality filtering and variant calling of raw sequencing data used a modified Stacks v1.3 pipeline (Catchen et al. 2013, Andrews 2018). We required a minimum read depth of six and presence in at least three individuals to call a locus, following the recommendations of Mastretta-Yanes et al. (2015) for minimizing error rates while maximizing true biological variation. Using Bowtie (Langmead et al. 2009), we removed reads that aligned to Huang et al.’s (2014) *P. fremontii* chloroplast and Kersten et al.’s *P. tremula* x *P. alba* (2016) mitochondrial reference genomes (NCBI accessions NC_024734.1 and NC_028329.1). The final dataset consisted of 322 *P. fremontii* genotypes, represented by 8,637 SNPs filtered to one random SNP per locus.

### Endophyte genetic data

Total DNA was extracted from twig samples with DNeasy Plant Mini Kits (Qiagen). Fungal ITS2 rDNA was selectively amplified with 1 μM fungal-specific primers 5.8SFun and ITS4Fun (Taylor et al. 2016) using Phusion Green Hotstart II High-Fidelity PCR Master Mix (Thermo Scientific). Amplification and indexing were performed following Alvarado et al. (2018). Libraries were pooled and sequenced on a MiSeq Desktop Sequencer in 2×300 mode.

Demultiplexing and quality filtering were performed with the `split_libraries_fastq.py` command in QIIME 1.9.1 (Caporaso et al. 2010) using a minimum quality threshold of q20 and 0 bad characters; only reads which satisfied these requirements for ≥95% of their length were retained. Chimeras were removed using the – `uchime_ref` option in VSEARCH 1.1.1 (Rognes et al. 2016) against the UNITE chimera reference (Nilsson et al. 2015) for fungi. Sequences were de-replicated on the first 100 bases using QIIME’s prefix/suffix OTU picker. OTU picking was performed *de novo* with Swarm (Mahé et al. 2014) at d4 resolution (~98.2% similarity). Taxonomic identities were assigned with BLAST in QIIME (maximum e-value = 0.001, 90% minimum sequence identity) against the dynamic UNITE database (Kõljalg et al. 2013). OTUs constituting <0.005% of the total dataset were removed (Bokulich et al. 2013). OTU tables were rarefied to the lowest sample depth for the purpose of assessing alpha diversity or normalized with cumulative sum scaling (Paulson et al. 2013) for all other analyses.

### Populus fremontii genetic structure and phylogenetic relationships

Genetic clustering was first assessed with principal component analysis (PCA) using SNPRelate (Zheng et al. 2012) in R 4.0.2 (R Development Core Team 2020); three extreme outliers were identified and removed. We then applied the Bayesian clustering algorithm implemented in Structure 2.3.4 (Pritchard et al. 2000, Falush et al. 2003). Assuming admixture and an independent alleles model, we ran 60,000 Markov Chain Monte Carlo (MCMC) generations with a 20,000 generation burn-in for *K* = 1-10 populations. Six iterations were run for each *K*. We identified the best-supported *K* following the Evanno et al. (2005) method (Δ *K* statistic) implemented in Structure Harvester (Earl and vonHoldt 2012), then used CLUMPP 1.1.2 (Jakobsson and Rosenberg 2007) to merge replicate runs and generate a six-run consensus for the best-supported *K*. Results were visualized with Distruct 1.1 (Rosenberg 2004). Evolutionary relationships among *P. fremontii* sampling locations were estimated with a maximum-likelihood tree and GTR + Γ nucleotide evolution model in PhyML 3.0 (Guindon et al. 2010).

### Environmental data

We identified a suite of 20 environmental predictor variables that we hypothesized to be related to gene flow and connectivity among *P. fremontii* and its associated communities (Cushman et al. 2014; Table S2). Wind is an important dispersal mechanism for cottonwood pollen and seed, therefore we included average wind velocity vectors to test for the influence of directional resistance to prevailing spring (February-May) winds. Mean monthly wind data was derived from the NCEP North American Regional Reanalysis and averaged across 1979-1989 (Mesinger et al. 2006, NCEP 2017). *Populus fremontii* is an obligate riparian species, and gene flow in this species is restricted by terrestrial uplands and low order streams, whereas mid-sized to larger, higher order rivers facilitate gene flow (Cushman et al. 2014). Thus we included topographic wetness index (TWI), a continuous hydrological measure that incorporates both slope and upland drainage area in the calculation of steady-state wetness (Boehner et al. 2002), to account for the influence of hydrology on gene flow.

Cottonwood pollen dispersal and seedling establishment are also intimately linked to climate through its cueing of reproductive phenology and influence on timing of spring flood events (Rood et al. 2005, Cooper et al., 2019). Cushman et al. (2014) found that genetic differentiation increased with cumulative differences in winter and spring precipitation, and Ikeda et al. (2017) linked differentiation among *P. fremontii* ecotypes to variation in minimum winter temperature and precipitation seasonality. We included nine bioclimatic variables representing temperature and precipitation means, minimums, maximums, and seasonality. Data represent 30-year averages (1970-2000, WorldClim v2 (Fick and Hijmans 2017)). We predicted that arthropod and endophyte communities would also be strongly influenced by seasonal climate cues as their life cycles closely track phenology of resource availability in their hosts. For example, successful larval development requires alignment of hatch times with bud flush and leaf development, which vary by >55 days across *P. fremontii’s* range (Cooper et al. 2019).

### Identifying drivers of community similarity

To test for the influence of climate in driving genetic differentiation in *P. fremontii*, we conducted a PCA followed by a permutational multivariate analysis of variance (perMANOVA, 999 permutations, *adonis* function, vegan R package (Oksanen et al. 2019)). Differentiation in functional traits and associated communities among *P. fremontii* ecotypes was assessed using non-metric multidimensional scaling (NMDS), followed by analysis of similarity (ANOSIM) and multi-response permutation procedure (MRPP) in PC-ORD, respectively (McCune and Mefford 2016).

To assess the relative contributions of host tree genetics, geography, and environment in driving community assembly, we used partial Mantel tests (999 permutations). To better understand the scale at which host tree genetics influences community organization, we investigated tree genotype-community relationships via an individual-based model (considering all tree genotypes individually), and a population-based model (allele frequencies summarized per sampling location). Pairwise genetic distance among *P. fremontii* individuals was calculated using principal component analysis (PCA)-based genetic distance (Shirk et al. 2010, Bothwell et al. 2017) on the first 15 PC axes, using SNPRelate and ecodist R packages (Zheng et al. 2012, Goslee and Urban 2007). Genetic differentiation (*F*_ST_) among sampling locations was derived from Stacks (Catchen et al. 2013). Individual-based and site-level community relatedness matrices were constructed using Bray-Curtis dissimilarity, incorporating both composition and abundance data (Bray and Curtis 1957). We used redundancy analysis (RDA) to identify which environmental variables had the strongest influence on macroscale community assembly.

## Results

### Population genetic structure and phylogenetic relationships

PCA (Fig. S1) and Structure (Fig. 1) analyses revealed that *P. fremontii* is differentiated into three primary genetic groups (*K* = 3). Our dataset including 8,637 SNPs is largely in agreement with the pattern of genetic structure found by Cushman et al. (2014), using 13 MSAT markers. Based on Cushman et al.’s (2014) genetic groupings, Ikeda et al. (2017) defined three *P. fremontii* ecotypes within the coterminous US, which occupy significantly different climate niches: Sonoran Desert (SD), Central California Valley (CCV), and Utah High Plateau (UHP). We extended sampling efforts to include the full species’ range, with additional sampling targeting southern California, northern Utah, and the Méxican states of Sonora and Durango. Interestingly, the more extensive collections did not reveal additional primary genetic structure; *K* = 3 remained the best supported number of populations. However, both PCA and an additional Δ*K* peak at *K* =6 indicated support for secondary hierarchical substructuring (Fig. S1). Therefore, we ran additional Structure analyses within each of the three primary genetic groups, following the same methods as above. Hierarchical substructure analysis supported *K* = 6, with each of the three ecotypes further split in two (Fig. S2). Based on these findings, we amend Ikeda et al.’s (2017) ecotype designations to separate Southern California (SC) from the more northern CCV ecotype, split the SD ecotype into Northern Sonoran Desert (NSD) and Southern Sonoran Desert (SSD) ecotypes, and differentiate the Utah High Plateau (UHP) from the Southern Colorado Plateau (SCP; Fig. 2). Phylogenetic relatedness among *P. fremontii* sampling locations further supports genetic divergence with strong branch support (95-100%) for the three primary ecotypes (Fig. 3). Additional substructure nests monophyletic groups by geography within the three major clades.

**Figure 1.**
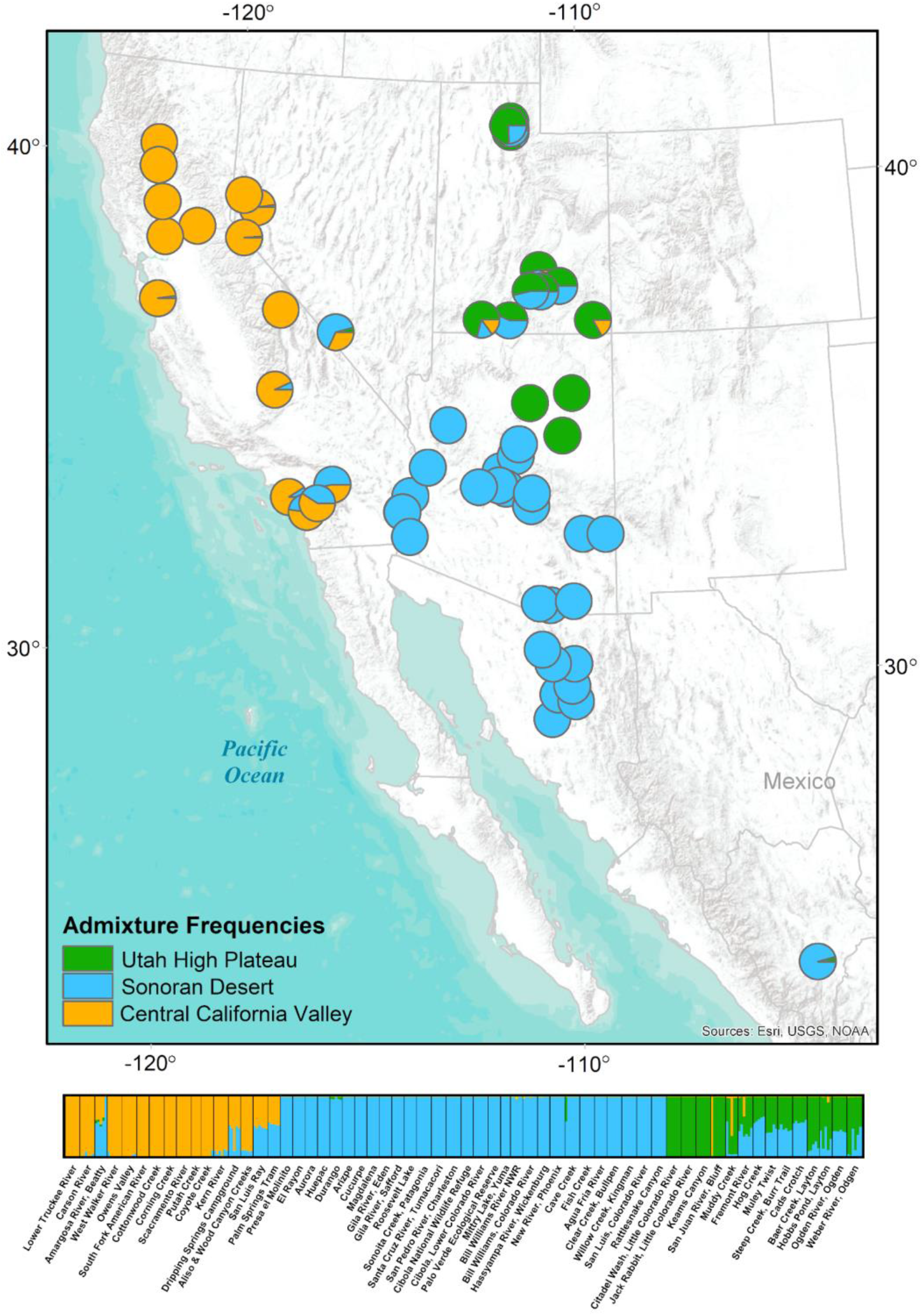
*Populus fremontii* segregates into three primary genetic lineages across its range. Different colored pies and bars in the genetic structure analysis *q* plot indicate admixture frequencies among genetic groups.

**Figure 2.**
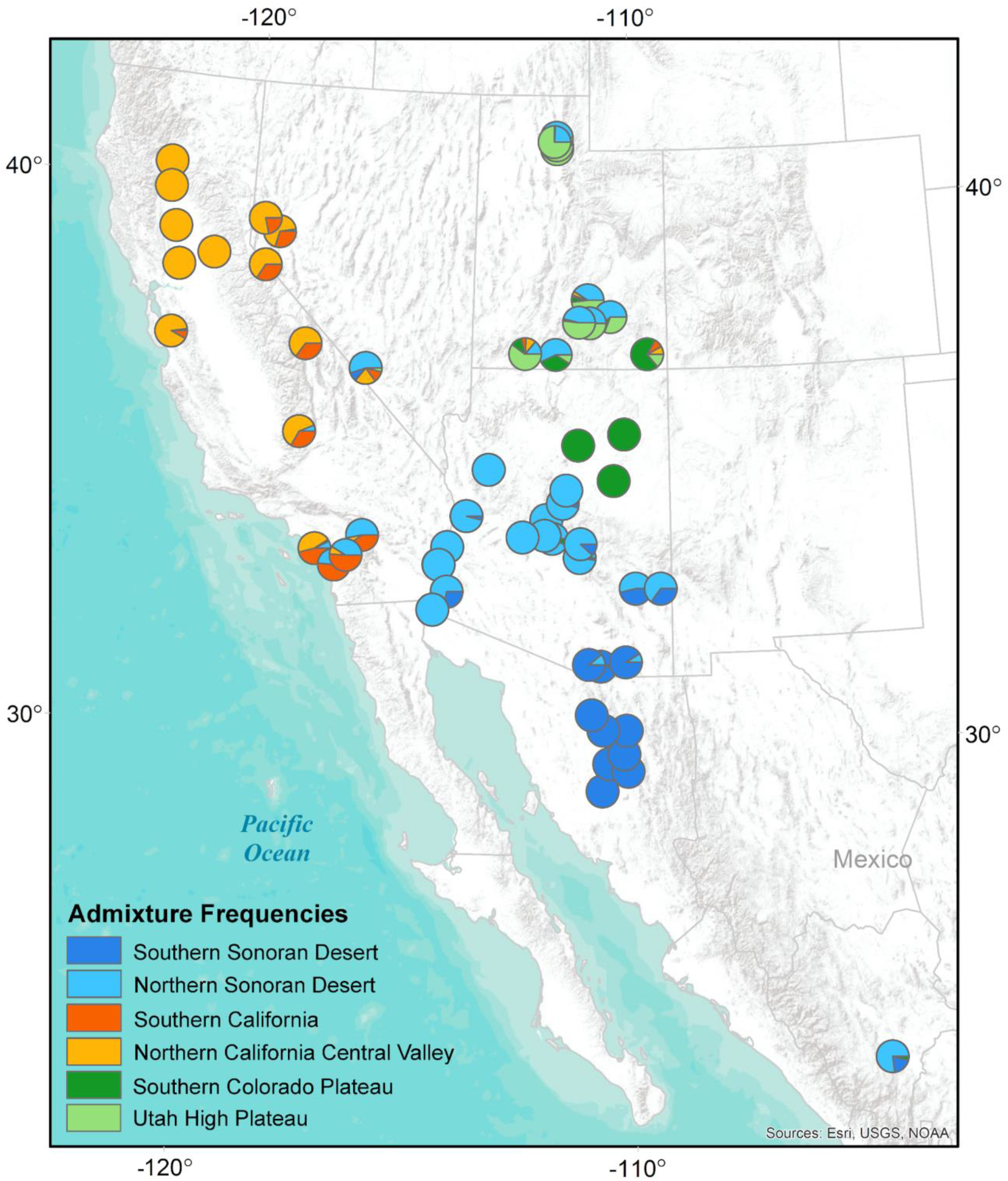
Admixture frequencies illustrate secondary hierarchical structure (*K* = 6) within the three primary ecotypes.

**Figure 3.**
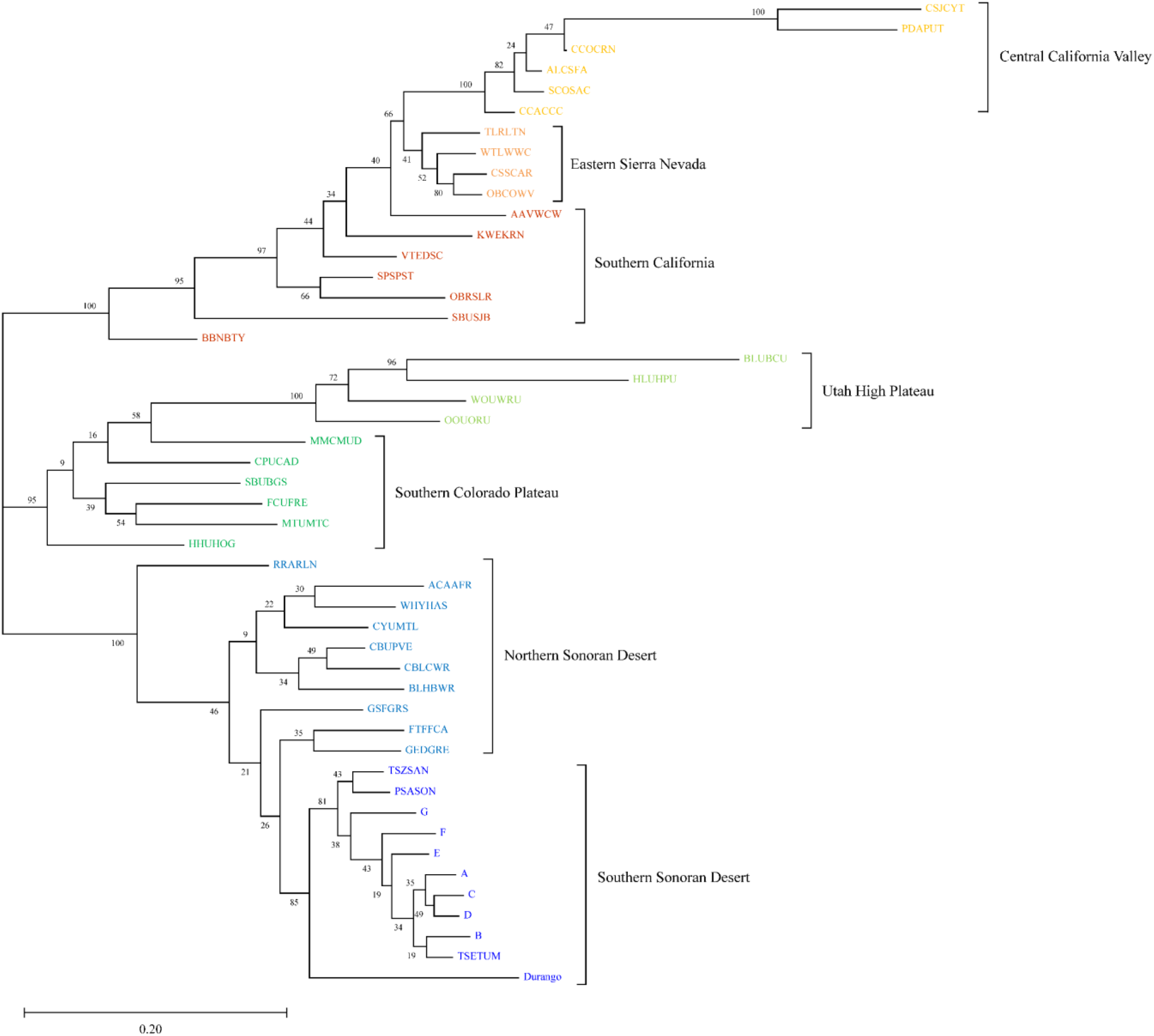
Phylogenetic relationships within *P. fremontii* show strong support for three primary genetic lineages and monophyletic clustering by ecoregion.

### Identifying drivers of community similarity

We predicted that *P. fremontii* genetic structure would be associated with regional variation in environmental selection pressures, and that more similar host tree genotypes would support more similar associated community phenotypes. We first tested whether *P. fremontii* population genetic groups occupy distinct climate niches, and found that 54.5% and 55.1% of the variation in climate space could be attributed to *K* = 3 and *K* = 6 genetic groups, respectively (perMANOVA *R*^2^ = 0.55, *p* < 0.001). The first two PC axes explained 59.7% of variation among ecotypes (Fig. 4), and revealed several notable patterns of niche divergence. The Utah High Plateau (UHP) ecotype is characterized by colder winters, the greatest temperature seasonality, and the highest dry quarter precipitation. In contrast, the Central California Valley (CCV) ecotype experiences the greatest precipitation seasonality, the highest winter precipitation, and the greatest summer heat moisture index (i.e., both high heat and low moisture contribute to high aridity). The Sonoran Desert (SD) ecotype’s climate niche is moderate with respect to precipitation seasonality, with the summer monsoon from the Gulf of México and winter storms from the Pacific bringing roughly equal contributions of annual precipitation. This ecotype also experiences the hottest summer temperatures, longest growing season, and greatest annual climate moisture deficit (i.e., largest sum of monthly differences between evaporation and precipitation). While differences in climatic selection pressures may influence host genetic divergence, associated communities ultimately cue into functional traits of the host. Accordingly, genetic and climate niche differentiation were also associated with differentiation in plant growth characteristics. Number of leaves, leaf area, twig diameter, and twig length associated with 3year growth collectively showed significant differentiation among ecotypes (ANOSIM *R* = 0.151, *p* = 0.01). Supporting our prediction, host tree ecotype emerged as a significant predictor of community phenotype for both arthropods and endophytes (Fig. 5, MRPP, *p* < 0.0001, *p* = 0.04, respectively).

**Figure 4.**
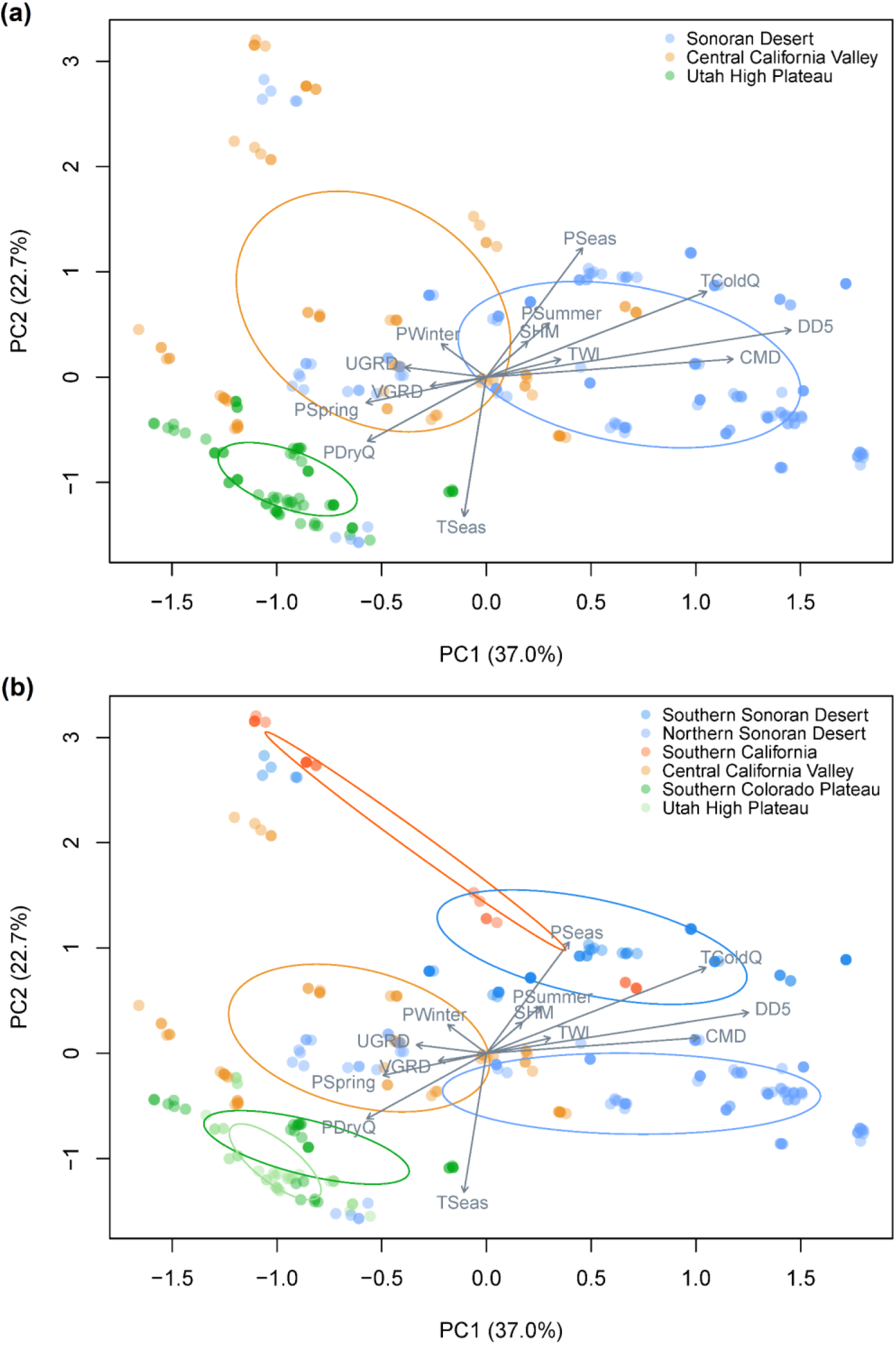
*Populus fremontii* ecotypes occupy significantly different climate niches, with population genetic group explaining (a) 54.5% and (b) 55.1% of the variation in climate space for *K* = 3 and *K* = 6, respectively (*p* = 0.001).

**Figure 5.**
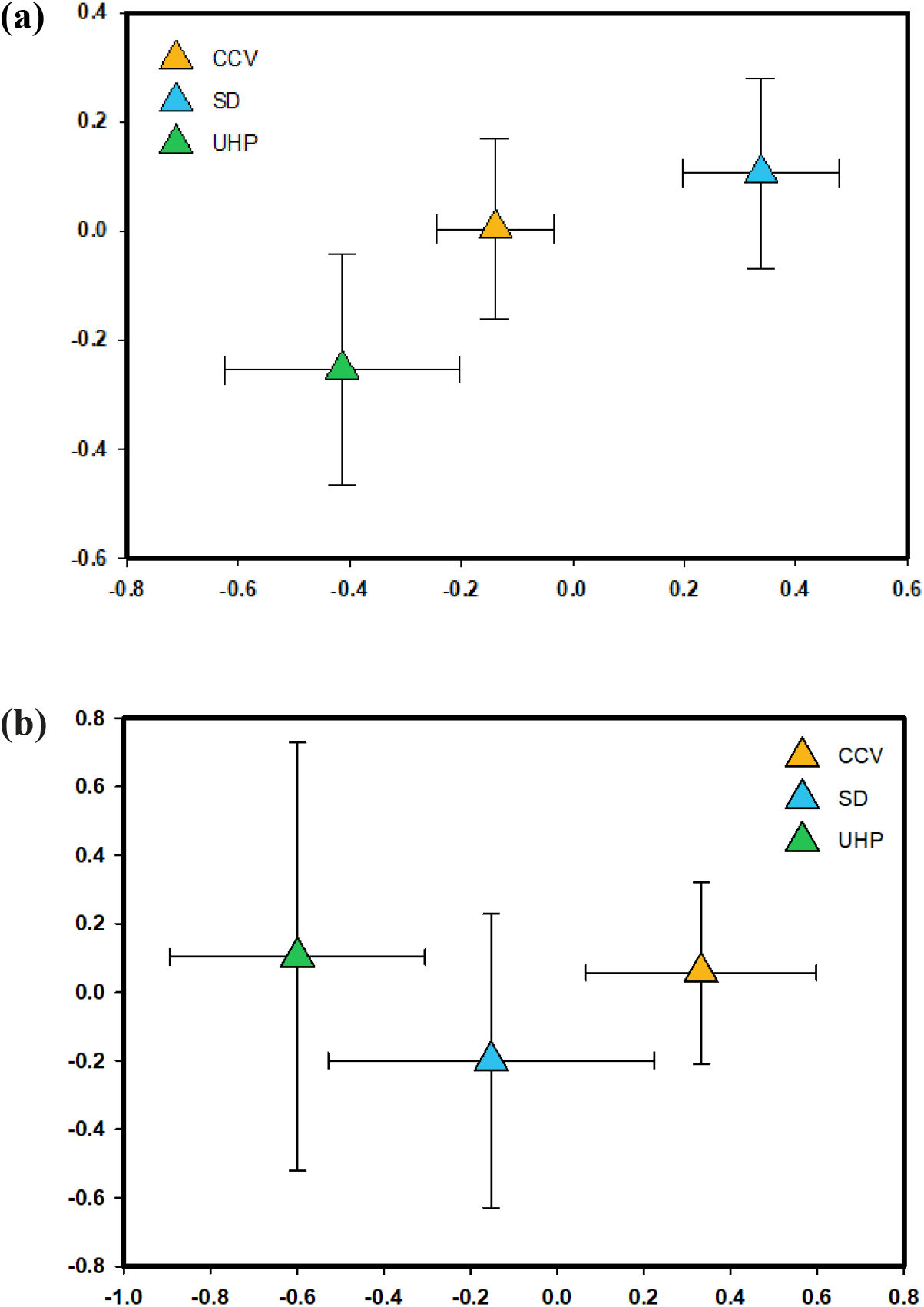
(a) Leaf modifying arthropod and (b) twig fungal endophyte communities are significantly differentiated across ecotypes (non-metric multidimensional scaling (NMDS, stress score = 0.19) followed by a multi-response permutation procedure (MRPP, *p* < 0.0001, *p* = 0.04, respectively).

We next quantified the relative contributions of host tree genetics, geography, and environment in driving community assembly, using a multi-scale approach which considered both individual- and stand-level relationships (Table 1). For the fungal endophyte community, geographic and environmental distance contributed equally to community assembly across individual host genotypes (partial Mantel *r* = 0.09, *p* = 0.07, *p* = 0.08, respectively), however these relationships were marginally significant, and we detected no independent effect of genetic distance. At the scale of forest patches, environmental distance was no longer significant. Instead genetic distance emerged as the primary driver, explaining 17% of variation in community similarity (*p* = 0.04), with a smaller contribution from geographic distance (15%, *p* = 0.03). However, reciprocal partial Mantel tests partitioning out the effects of genetic and geographic distance on each other were non-significant. Given this confounding result, we tested for isolation by distance (IBD). We detected a weak relationship between genetic and geographic distance among individual tree genotypes (Mantel *r* = 0.095, *p* = 0.02), however IBD was substantial at the scale of forest patches (Mantel *r* = 0.477, *p* = 0.001).

**Table 1.** Mantel *r* correlations quantifying the relative contribution of genetic (GenD), geographic (Geog), and environmental distance (EVD) in driving community similarity at the scale of (a) individual tree genotypes and (b) forest patches. Simple Mantel tests are represented along the diagonal (left to right). Remaining values represent partial Mantel tests, with the amount of variation in the associated community being explained by the column variable while partitioning out the effect of the row variable. For example, 9% of the variation in the endophyte community across individual tree genotypes can be explained by geographic distance after accounting for the effect of environmental distance, and this relationship is marginally significant.

| ENDOPHYTES |  |  |  | ARTHROPODS |  |  |  |
| --- | --- | --- | --- | --- | --- | --- | --- |
|  | Geog | EVD | GenD |  | Geog | EVD | GenD |
| Geog | 0.16** | 0.09† | 0.07 | Geog | 0.08** | 0.17*** | -0.03 |
| EVD | 0.09† | 0.16** | 0.04 | EVD | -0.02 | 0.18*** | -0.05 |
| GenD | 0.15** | 0.14* | 0.09 | GenD | 0.08** | 0.19*** | -0.02 |

(b)
|  | Geog | EVD | GenD |  | Geog | EVD | GenD |
| --- | --- | --- | --- | --- | --- | --- | --- |
| Geog | 0.15* | 0.01 | 0.1 | Geog | 0.44*** | 0.24* | -0.09 |
| EVD | 0.12† | 0.09 | 0.16† | EVD | 0.15† | 0.47*** | -0.16 |
| GenD | 0.05 | 0.07 | 0.17* | GenD | 0.43*** | 0.48*** | 0.13† |

$p < 0.1$ † $p < 0.05$ \* $p < 0.01$ \*\* $p < 0.001$ \*\*\*

For leaf modifying arthropods, environmental distance was the primary driver of community assembly, accounting for 17% (*p* = 0.001) of variation among individual trees after partialling out the effect of geographic distance. No independent geographic effect was observed after accounting for environmental distance. This relationship was amplified at the scale of forest patches, with environment independently explaining 24% (*p* = 0.02) of the variation in community assembly, and a smaller independent contribution detected for geography (Mantel *r* = 0.15, *p* = 0.06). At the individual tree scale, genetic distance was not significant (Mantel *r* = −0.024, *p* = 0.66). Conversely, host genetics explained 13% of community similarity among forest patches (*p* = 0.06), although this relationship was confounded with the effects of geography and environment.

We further investigated which environmental variables were most important in structuring leaf modifying arthropods across southwestern North America. Similar to the forest patch scale, environment explained 24.8% of total variation across the whole macrosystem (i.e., all *P. fremontii* ecotypes; Fig. S3, *p* = 0.001). While many species were generalists present across all ecotypes, several species emerged as indicators most commonly associated with specific ecoregions. *Pemphigus populicaulis* was common throughout southern California and found in limited numbers at two sites along the Arizona-California border, but nowhere else. *Pemphigus transversus* was most strongly associated with higher summer precipitation common to the SD ecoregion. Heliozelidae *Coptodisca* ssp. and *Coleophora* ssp. had limited presence across all three ecotypes, but were most abundant in association with the Sonoran Desert’s hotter temperatures and longer growing season. The cottonwood leafminer moth, *Paraleucoptera albella*, was most common throughout the Utah High Plateau, and to a lesser extent also in the Eastern Sierra Nevada.

## Discussion

Macroecological patterns arise from multiple abiotic and biotic processes interacting across local to continental scales. Here, we investigated the hypothesis that environmental variation, geography, and tree genetic connectivity jointly predict macroscale community assembly patterns across southwestern North America. As predicted, tree genetic connectivity was a significant factor contributing to assembly for both communities investigated, however we observed a closer association for fungal endophytes compared to leaf modifying arthropods. Tree genetics was the strongest predictor of macroscale endophyte assembly, whereas environment was a stronger predictor for arthropods. By investigating community assembly within a genetic framework, we demonstrate the feasibility of merging macrosystems ecology with community genetics, an objective that we consider critical for predicting global change impacts on macrosystem properties (e.g., widespread biodiversity loss).

### Climate as a driver of host tree local adaptation

Climate has long been known to be a primary driver of broadscale species distributions (Woodward 1987), and substantial evidence of local adaptation in response to climate has been observed in *P. fremontii* common gardens for traits related to growth, phenology, and temperature regulation (Grady et al. 2013, Fischer et al. 2017, Cooper et al. 2019, Hultine et al. 2020). The strong genetic divergence and environmental niche partitioning found here lend further support for local adaptation among ecotypes. While our current study targeted more extensive sampling efforts throughout southern California, northern Utah, and México, new samples nested within the original three ecotypes defined by Ikeda et al. (2017). Yet, local adaptation can occur within ecotypes at finer scales of even a few hundred meters or less (Urban et al. 2020). Investigating hierarchical substructure within each of the three primary ecotypes revealed additional partitioning, with substructure generally segregating along latitudinal gradients into northern and southern populations within each primary ecotype.

We detected significant niche divergence when considering both *K* = 3 and 6 genetic groups. Greater overlap was observed between CCV and SD climate niches, relative to UHP (Fig. 4a). The UHP ecotype occupied a substantially narrower and more divergent climate niche relative to the other ecotypes, even though samples spanned comparable latitudinal ranges. While both Structure analysis and phylogenetic distance indicated strong neutral differentiation between northern and southern UHP populations, climate niche differentiation between populations within this lineage was much less than that observed within the SD and CCV ecotypes (Fig. 4b). In contrast, all four substructure groups within the latter ecotypes exhibited significant niche divergence. Previous studies coupled with the strong divergence observed here among climate niches and leaf and twig traits suggest that ecotypes of *P. fremontii* are locally adapted. We therefore predicted that macroscale patterns of associated community assembly would reflect patterns of genetic connectivity of the host tree, and ultimately functional trait diversity among *P. fremontii* ecotypes.

### Predicting macroscale community assembly

We investigated the relative contributions of host tree genetics, geography, and environment in driving community assembly for two closely associated communities: twig fungal endophytes and leaf modifying arthropods. Contrary to our prediction, we detected no significant relationship with tree genetic connectivity for either community at the individual tree scale. Cushman et al. (2014) found that the majority of genetic diversity is distributed within individuals in this species (75%), with only 3% distributed among individuals and 22% among populations. Given the very low diversity distributed among individuals, it is perhaps not surprising that a signal of host genetic connectivity was undetectable for the individual-based model. Common garden studies have consistently observed that plant genotype predicts heritable community phenotypes in this species (Wimp et al. 2005, Schweitzer et al. 2008, Keith et al. 2010, Ferrier et al. 2012, Lamit et al. 2015). For example, host genotype was found to explain 33% of variation in fungal pathogen community structure on *P. fremontii* (Busby et al. 2013). At the macroscale, environmental variation and geographic distance instead overwhelmed the signal of genetic connectivity among individual tree genotypes.

When considering genetic connectivity among forest patches, however, host-community relationships revealed spatial nonstationarity. At this scale, we observed positive correlations for both communities. As predicted, host genetic connectivity had a stronger influence on endophyte (17%) compared to the arthropod community assembly (13%), and in fact was the most important determinant of macroscale endophyte structure. In contrast, the independent effect of environmental variation explained nearly twice the variation (24%) in macroscale arthropod community assembly relative to host genetics (13%).

Community properties like diversity, stability, and structure are often viewed as emergent properties, because they result from complex interactions that cannot be easily explained by first principles (Herfeld & Ivanova 2020). However, Keith et al. (2010) experimentally showed in common garden studies that arthropod diversity and community stability across years were heritable community phenotypes; some tree genotypes innately supported richer or more depauperate communities that were more or less stable across years, respectively. Wimp et al. (2005) also found that different tree genotypes support different communities; thus, the greater the genetic diversity harbored in foundation species, the greater community biodiversity they can support. Such findings, including the results presented here, argue that community properties arise in part as a direct outcome of first genetic principles in which the genetic-based multivariate phenotypes of plants can be used to predict associated properties of dependent communities at multiple scales.

### Conservation management implications

Arthropods have been particularly hard-hit by the global extinction crisis (Thomas et al. 2004, Hallmann et al. 2017), with an estimated 40% of all insect species exhibiting precipitous declines (Sánchez-Bayo and Wyckhuys 2019). Macroscale patterns of microbial diversity are less well understood, but no less important for supporting ecosystem services that all higher orders depend on (Banerjee et al. 2020). Previous work has documented the disproportionately high biodiversity that *P. fremontii* riparian forests support relative to surrounding aridlands (Poff et al. 2012), and our findings highlight that tree genetic variation is a significant factor driving this biodiversity.

Conservation of regional biodiversity relies on preserving the remaining genetic variation harbored within this foundation species. *Populus fremontii* riparian forests are among the most threatened ecosystems in the US; as a result of water diversion, drought, and land-use changes, <3% of this species’ pre-20^th^ century distribution remains (Noss et al. 1995). Climate change is predicted to further reduce *P. fremontii* s suitable habitat, particularly for the UHP ecotype (Ikeda et al. 2017), which occupies the narrowest climate niche. Climate change will not only require species to migrate, adapt, or perish (Aitken et al. 2008), but it may also reduce tree productivity, which in turn alters productivity-diversity relationships and foundation species’ capacity to support diverse associated communities (Ikeda et al. 2014). Evans et al. (2016) used common gardens of the same *P. angustifolia* genotypes reciprocally transplanted across their natural range from Arizona to Alberta to investigate phenotypic differentiation in functional traits associated with latitude of origin, emphasizing the role climate has played in divergent selection. They found evidence of arthropod segregation across different tree populations within the common garden environment that were consistent with tree trait divergence, and in particular, community metrics were positively correlated with tree productivity. These results lend support to the hypothesis that divergence in associated communities tracks genetically-based differences in phenology and growth traits of host plants at macrosystem scales. While our findings show clear differentiation among endophytes and arthropods across ecotypes (Fig. 5), our results indicate that different communities and species within those communities may exhibit idiosyncratic and specialized responses to global change (Fig. S3).

We identified three primary and six secondary *P. fremontii* ecotypes, which exhibit genetic and functional trait divergence associated with unique climate niches, that in turn contribute to macroscale biodiversity patterns. Regional management plans should aim to maintain high genetic diversity within local forest patches and support conservation genetic management of representative heterogeneity across the six ecotypes. River network connectivity is positively correlated with genetic diversity in *Populus* ssp. (Bothwell et al. 2017), thus maintaining existing riparian corridors and restoring degraded stretches will be critical for conserving this foundation species and the diverse communities it supports.

## Notes

### Competing Interest Statement

The authors have declared no competing interest.

