## Supplementary Information for "Macrosystem community assembly patterns are predicted by foundation tree species genetic connectivity and environment across the American Southwest"

### Tables & Figures

**Table S1.** *Populus fremontii* collection information.

**Table S2.** Environmental predictor variables.

**Figure S1.** Principal component analysis of *Populus fremontii* genetic structure.

**Figure S2.** *Populus fremontii* admixture frequency *q* plots.

**Figure S3.** Arthropod differentiation by environment.

**Table S1.** *Populus fremontii* collection information. We sampled leaf material and recorded geographic coordinates for 453 individual trees at 58 sampling locations throughout the southwestern US and northwestern México for genomic analysis. Latitude and longitude represent site means (North American Datum 1983 coordinate system). GPS accuracy varied +/- 2 to 18 feet.

| Sampling location | Region | Site ID | Sample size (*n*) | Latitude | Longitude |
|---|---|---|---|---|---|
| Presa el Molinito | México | A | 5 | 29.209509 | -110.729978 |
| Aliso & Wood Canyon Creeks | CA | AAVWCW | 5 | 33.541433 | -117.736583 |
| Agua Fria River | AZ | ACAAFR | 5 | 34.236500 | -112.098567 |
| South Fork American River | CA | ALCSFA | 5 | 38.803178 | -120.909905 |
| El Rayon | México | B | 5 | 29.716004 | -110.576465 |
| Amargosa River, Beatty | NV | BBNBTY | 5 | 36.908835 | -116.754439 |
| Clear Creek, Bullpen | AZ | BCEBUL | 12 | 34.539720 | -111.696600 |
| Bill Williams River | AZ | BLHBWR | 5 | 34.276762 | -114.056059 |
| Baer Creek, Layton | UT | BLUBCU | 5 | 41.025100 | -111.918510 |
| Aurora | México | C | 5 | 29.574928 | -110.130366 |
| Cibola National Wildlife Refuge | AZ | CBLCWR | 5 | 33.363128 | -114.701780 |
| Palo Verde Ecological Reserve | AZ | CBUPVE | 5 | 33.695717 | -114.501597 |
| Cottonwood Creek | CA | CCACCC | 6 | 40.374864 | -122.283527 |
| Corning Creek | CA | CCOCRN | 5 | 39.925465 | -122.223639 |
| Cave Creek | AZ | CCUCAV | 7 | 33.890000 | -111.951000 |
| Citadel Wash, Little Colorado River | AZ | CLFLCR | 11 | 35.613000 | -111.319000 |
| Cads Crotch | UT | CPUCAD | 6 | 37.266779 | -111.906819 |
| Coyote Creek | CA | CSJCYT | 6 | 37.278826 | -121.804032 |
| Carson River | NV | CSSCAR | 6 | 39.287469 | -119.240370 |
| Mittry Lake, Yuma | AZ | CYUMTL | 4 | 32.869694 | -114.476672 |
| Huépac | México | D | 5 | 29.893426 | -110.221022 |
| Durango | México | Durango | 5 | 24.012614 | -104.492626 |
| Arizpe | México | E | 5 | 30.339434 | -110.161876 |
| Cucurpe | México | F | 5 | 30.340113 | -110.706744 |
| Fremont River | UT | FCUFRE | 6 | 38.274301 | -111.084869 |
| Fish Creek | AZ | FTFFCA | 6 | 33.538617 | -111.274333 |
| Magdalena | México | G | 5 | 30.625795 | -110.979590 |
| Gila River, Eden | AZ | GEDGRE | 5 | 32.972722 | -109.916698 |
| Gila River, Safford | AZ | GSFGRS | 5 | 32.965227 | -109.309788 |
| Hog Creek, Hog Springs Picnic Area | UT | HHUHOG | 5 | 37.961527 | -110.494506 |
| Hobbs Pond, Layton | UT | HLUHPU | 5 | 41.097770 | -111.926930 |
| Jack Rabbit, Little Colorado River | AZ | JLAJAK | 12 | 34.960000 | -110.436000 |
| Keams Canyon | AZ | KKHOPI | 10 | 35.811520 | -110.169580 |
| Kern River | CA | KWEKRN | 6 | 35.672660 | -118.327436 |
| Willow Creek, Kingman | AZ | KWFWIL | 11 | 35.143000 | -113.542840 |

| | | | | | |
|---|---|---|---|---|---|
| Muddy Creek | UT | MMCMUD | 5 | 37.277458 | -112.689254 |
| Rattlesnake Canyon | AZ | MRNRAT | 12 | 34.783050 | -111.613720 |
| Muley Twist | UT | MTUMTC | 5 | 37.848650 | -111.030573 |
| New River, Phoenix | AZ | NRVNEW | 16 | 33.947660 | -112.136170 |
| Owens Valley | CA | OBCOWV | 6 | 37.281940 | -118.334401 |
| San Luis Ray | CA | OBRSLR | 6 | 33.264917 | -117.233983 |
| Ogden River, Ogden | UT | OOUORU | 6 | 41.235900 | -111.933110 |
| Putah Creek | CA | PDAPUT | 6 | 38.527173 | -121.794660 |
| Sonoita Creek, Patagonia | AZ | PSASON | 17 | 31.533758 | -110.765454 |
| Roosevelt Lake | AZ | RRARLN | 5 | 33.795117 | -111.255367 |
| Steep Creek, The Gulch, Burr Trail | UT | SBUBGS | 6 | 37.860553 | -111.311855 |
| San Juan River, Bluff | UT | SBUSJB | 6 | 37.273264 | -109.556586 |
| Scacramento River | CA | SCOSAC | 6 | 39.208535 | -121.984810 |
| San Luis, Colorado River | AZ | SCTMEX | 13 | 32.527020 | -114.803690 |
| Palm Springs Tram | CA | SPSPST | 5 | 33.844250 | -116.597317 |
| Lower Truckee River | NV | TLRLTN | 6 | 39.509220 | -119.651311 |
| Santa Cruz River, Tumacacori | AZ | TSETUM | 18 | 31.565745 | -111.045160 |
| San Pedro River, Charleston | AZ | TSZSAN | 19 | 31.610062 | -110.167571 |
| Dripping Springs Campground | CA | VTEDSC | 5 | 33.453500 | -116.969317 |
| Hassyampa River, Wickenburg | AZ | WHYHAS | 17 | 33.906933 | -112.675167 |
| Weber River, Odgen | UT | WOUWRU | 6 | 41.157497 | -111.991251 |
| West Walker River | CA | WTLWWC | 6 | 38.660089 | -119.539252 |

| Wind | Description | Source |
|---|---|---|
| Mean Spring wind velocity U vectors<br>Mean Spring wind velocity V vectors | Wind vectors represent ten year averages (1979-1989) of February-May monthly means, measured 10m aboveground. | Mesinger et al. (2006); NCEP North American Regional Reanalysis (NARR); https://rda.ucar.edu/datasets/ds608.0/ |
| **WorldClim v2** | | |
| Temperature seasonality (BIO4)<br>Maximum temperature of the warmest month (BIO5)<br>Minimum temperature of the coldest month (BIO6)<br>Mean temperature of the coldest quarter (BIO11)<br>Mean annual precipitation (BIO12)<br>Mean precipitation of the driest month (BIO14)<br>Precipitation seasonality (CV) (BIO15)<br>Mean precipitation of the driest quarter (BIO17)<br>Mean precipitation of the coldest quarter (BIO19) | Bioclimatic variables represent thirty year averages (1970-2000) of means, maximums, minimums, and variation. | Fick & Hijmans (2017); http://worldclim.org/version2 |
| **ClimateNA** | | |
| Hargreave's climatic moisture index (CMD)<br>Degree-days above 5°C (DD5)<br>Summer precipitation (PPT_SM; June-August)<br>Spring precipitation (PPT_SP; March-May)<br>Winter precipitation (PPT_WT; December-February)<br>Summer heat moisture index (SHM; mean warmest month temperature/(mean summer precip./1000))<br>Continentality (TD; difference between mean coldest month and mean warmest month temperatures; °C) | Variables represent thirty year averages for the years 1981-2010. | Hamann et al. (2013), AdaptWest Project (2015); https://adaptwest.databasin.org |
| **Topographic Indices** | | |
| Topographic Wetness Index (TWI) | Derived from USGS SRTM DEM (2004) | Boehner et al. (2002); calculated in SAGA GIS |
| Heat Load Index (HLI) | | McCune et al. (2002) |

**Table S2.** Predictor variables, including prevailing Spring wind direction, bioclimatic, and topographic variables.

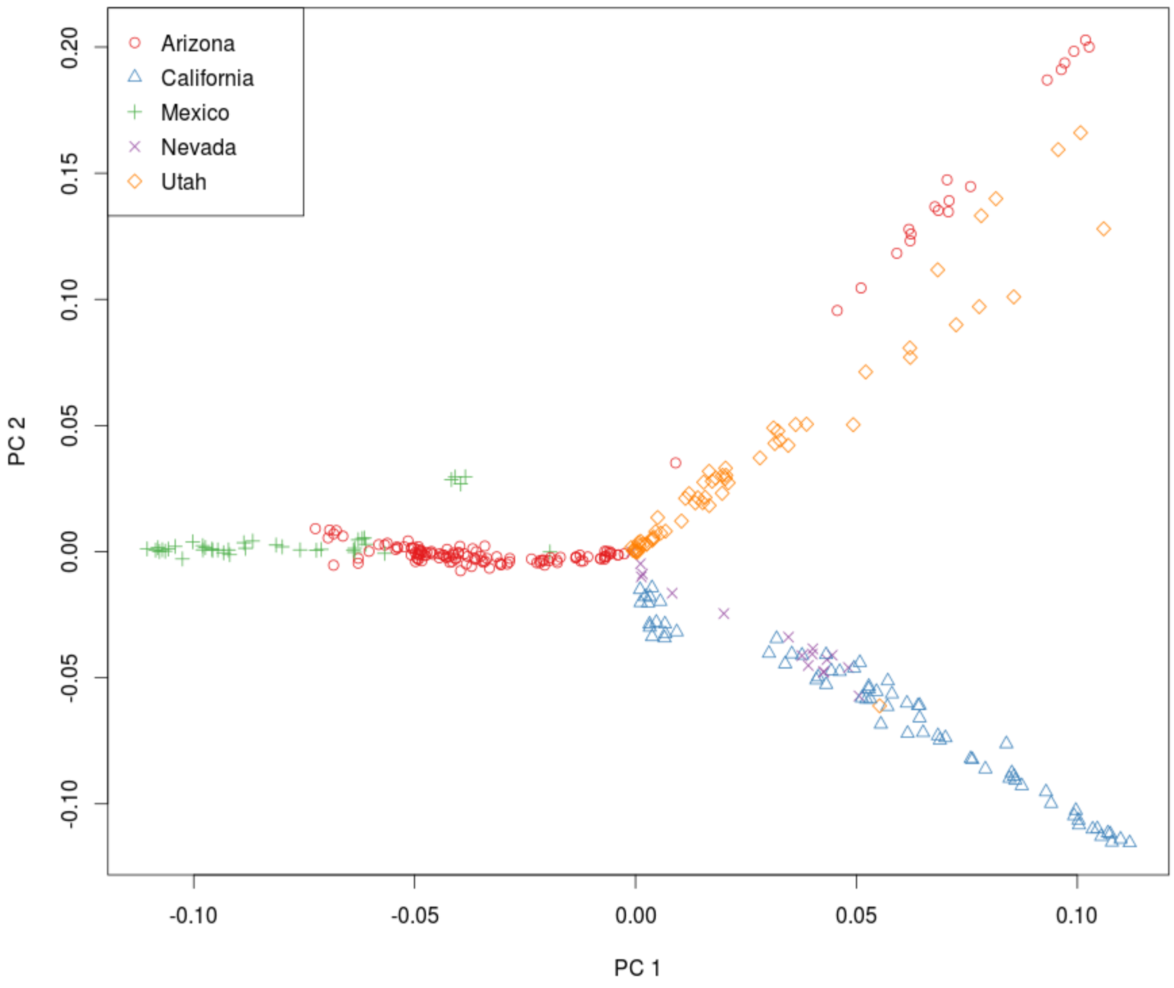


**Figure S1.** Principal component analysis (PCA) of *P. fremontii* genetic structure. The first 32 eigenvectors explained 40.15% of total variance (PC1 = 5%, PC2 = 4.5%). Data were subset to one random snp per locus, resulting in 8,637 snps across 322 samples.

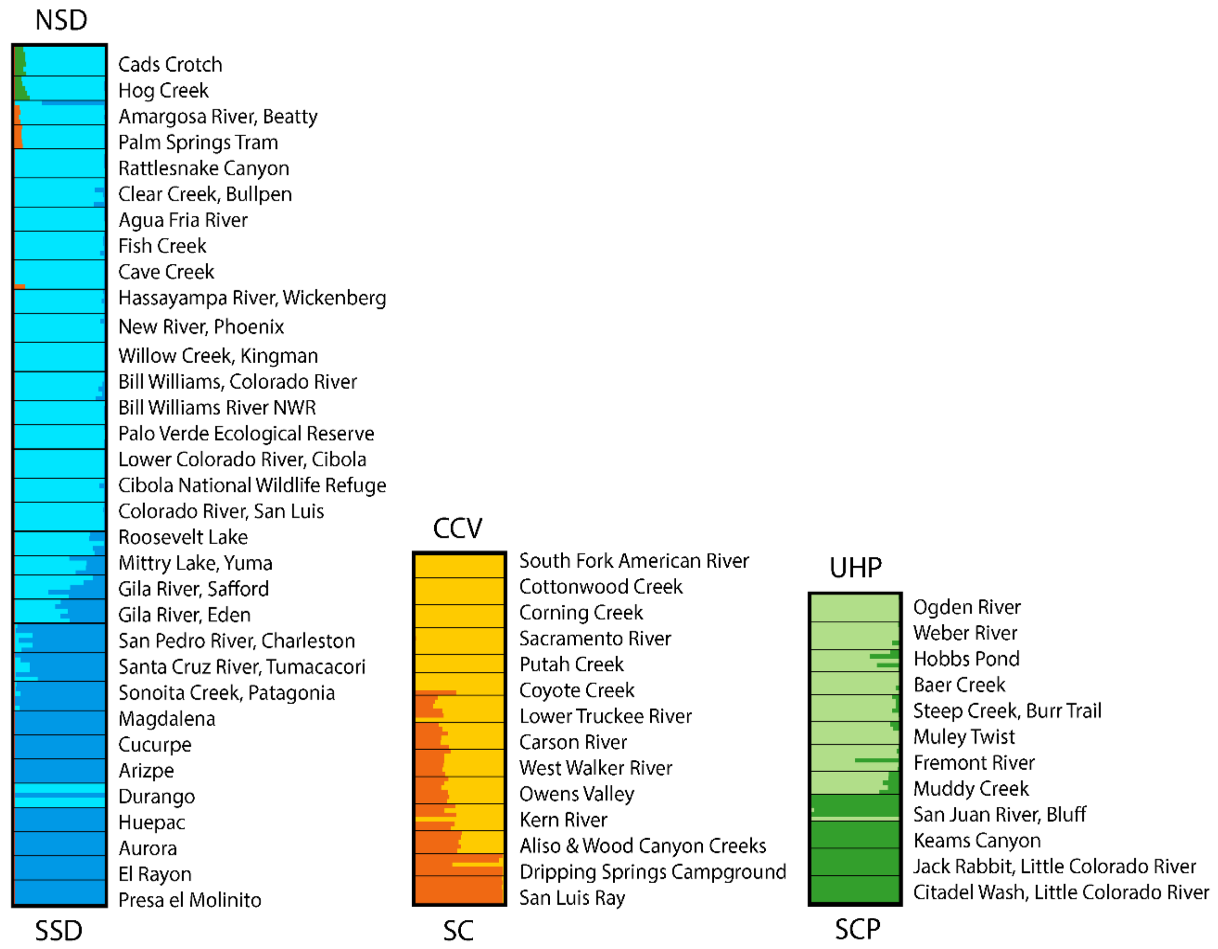


**Figure S2.** Hierarchical analysis of population genetic structure revealed secondary differentiation within each of the three primary ecotypes. NSD = Northern Sonoran Desert, SSD = Southern Sonoran Desert, CCV = Central California Valley, SC = Southern California, UHP = Utah High Plateau, SCP = Southern Colorado Plateau.

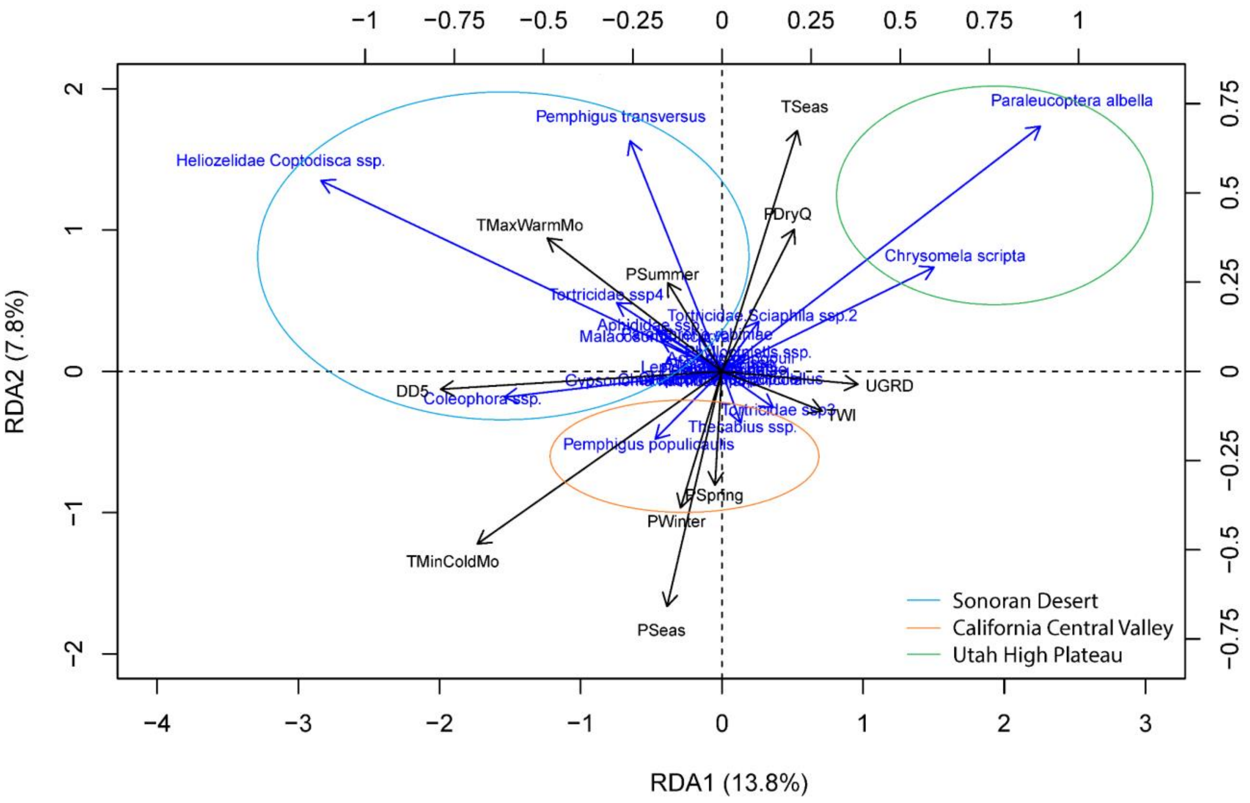


**Figure S3.** Twenty-five percent of the variation in the leaf modifying arthropod community is explained as a function of environment (RDA $R^2_{adj} = 0.248$, $p = 0.001$). Many species appear to be generalists present across all ecotypes, however several species emerged as specialists uniquely associated with specific ecotypes.
